# Acute hyperoxia improves spinal cord oxygenation and circulatory function following cervical spinal cord injury in the rat

**DOI:** 10.1101/2023.08.29.555438

**Authors:** Yen-Ting Lin, Kun-Ze Lee

## Abstract

Spinal cord injury is associated with spinal vascular disruptions that result in spinal ischemia and tissue hypoxia. This study evaluated the therapeutic efficacy of normobaric hyperoxia on spinal cord oxygenation and circulatory function at the acute stage of cervical spinal cord injury. Adult male Sprague–Dawley rats underwent dorsal cervical laminectomy or cervical spinal cord contusion. At 1–2 days after spinal surgery, spinal cord oxygenation was monitored in anesthetized and spontaneously breathing rats via the optical recording of oxygen sensor foils placed on the cervical spinal cord and pulse oximetry. The arterial blood pressure, heart rate, blood gases, and peripheral oxyhemoglobin saturation were also measured under hyperoxic (50% O_2_) and normoxic (21% O_2_) breathing. The results showed that contused animals had a significantly lower spinal cord oxygenation level than uninjured animals during normoxia. Cervical spinal cord contusion also significantly reduced peripheral oxyhemoglobin saturation, arterial oxygen partial pressure, and mean arterial blood pressure. Notably, the spinal oxygenation of contused rats could be improved to a level comparable to uninjured animals under hyperoxia. Furthermore, acute hyperoxia could elevate blood pressure, arterial oxygen partial pressure, and peripheral oxyhemoglobin saturation. These results suggest that normobaric hyperoxia can significantly improve spinal cord oxygenation and circulatory function during acute cervical spinal cord injury. We propose that adjuvant normobaric hyperoxia combined with other hemodynamic optimization strategies may prevent secondary damage after spinal cord injury and improve functional recovery.

## Introduction

Spinal cord injury damages neural cells and pathways and disrupts spinal cord microvascular structure and function. Figley et al. showed significantly reduced spinal vessel number and functional vasculature at one day post-injury in thoracic spinal cord contused rats (9). The dorsal spinal cord section also showed spinal tissue necrosis and decreased blood vessels in rats with thoracic spinal cord injury (7). These anatomical changes in spinal cord tissues subsequently impact spinal hemodynamics. Specifically, in a porcine model, spinal cord blood flow was significantly reduced during compression injury and remained lower than its pre-injury level even at 3 hours post-decompression (35). Li et al. also reported that spinal cord blood flow below the injury site was ∼50% lower than in the uninjured spinal cord in chronically injured rats (22). Hypoperfusion of the spinal cord results in spinal cord hypoxia, as reflected in a significantly reduced spinal cord oxygenation level (22, 35). These spinal pathological conditions will cause secondary injury cascades around the lesion epicenter and are unfavorable for the repair or regeneration of spared neural tissues (23). Therefore, several studies have suggested that preventing secondary injury may attenuate the progression of spinal pathology and improve recovery after spinal cord injury (8, 12, 40).

The therapeutic efficacy of hyperbaric oxygen treatment has been evaluated in animal and clinical studies (29). For example, hyperbaric oxygen treatment improved the locomotor activity of rats with thoracic spinal cord injuries (14, 36). Yaman et al. also showed that hyperbaric oxygen therapy could promote motor strength recovery after thoracic spinal cord injury in rats (44). Moreover, Smuder et al. reported that hyperbaric oxygen therapy could preserve diaphragm muscle specific force in cervically contused rats (33). These results suggest that hyperbaric oxygen therapy is a promising strategy for promoting functional recovery after spinal cord injury. However, hyperbaric oxygen therapy devices are not easily accessible and affordable in all hospitals. Moreover, hyperbaric oxygen treatment is associated with side effects, including barotrauma and central and pulmonary oxygen toxicity (13, 26). Therefore, several studies have used normobaric hyperoxia as an alternative approach, showing that it is beneficial after traumatic brain injury and stroke (3, 5, 15). However, whether normobaric hyperoxia is beneficial after cervical spinal cord injury remains unclear.

Cervical spinal cord injuries usually cause cardiorespiratory dysfunctions and result in systemic and central hypoxia due to damage to supraspinal respiratory and vasomotor pathways (10, 18, 19, 24, 27, 38, 41, 43). In addition, the cervical spinal cord has more extensive microvascular structures than other spinal cord segments (39). Consequently, injury to the cervical spinal cord segment may induce more severe hemorrhage and spinal cord tissue hypoxia, which may reduce hyperoxia’s efficacy. Therefore, this study examined whether acute normobaric hyperoxia treatment could effectively maintain tissue oxygenation in the cervical spinal cord and improve circulatory function in rats with mid-cervical spinal cord contusions.

## Materials and Methods

### Animals

This study used 28 male Sprague–Dawley rats aged 7–8 weeks purchased from BioLASCO Taiwan Co., Ltd. The animals were divided into an uninjured group (i.e., C3 laminectomy only; *n* = 12) and a contused group (i.e., C3 contusion; *n* = 16). All experimental protocols were approved by the Institutional Animal Care and Use Committee at National Sun Yat-sen University (approval number: 11001).

### Cervical spinal cord injury

Cervical spinal cord injury was induced at 9–10 weeks of age, as previously described (20, 21). The animal was anesthetized with xylazine (subcutaneous [s.c.], 10 mg/kg of Rompun; Bayer) and ketamine (intraperitoneal [i.p.], 140 mg/kg of Ketalar; Pfizer). In both uninjured and contused groups, the third cervical vertebra was fixed using two transverse clamps (#51692; Stoelting Co.) and Cunningham spinal adaptors (#51695; Stoelting Co.) mounted on a stereotaxic device followed by a dorsal C3 laminectomy. A contusion injury was induced in contused rats by releasing an impact rod (diameter: 2 mm; weight: 10 g; the Multicenter Animal Spinal Cord Injury Study impactor system) from a height of 6.25 mm above the midline C3 spinal cord. The dorsal muscle and skin were sutured with 4-0 sutures followed by injections of yohimbine (1.2 mg/kg, s.c.; Tocris), lactated Ringer’s solution (5 mL, s.c.; Nang Kuang Pharmaceutical Co., Ltd.), and buprenorphine (0.03 mg/kg, s.c.; Reckitt Benckiser Healthcare). Animals were given daily oral supplementation with Nutri-Plus Gel (1–3 mL; Vibac) and injections of lactated Ringer’s solution (5 mL, s.c.) until they resumed volitional eating and drinking. Two animals in the contused group died after contusion injury; these animals were excluded from the following experimental protocol.

### Measurement of physiological parameters

The animal was anesthetized with urethane (1.6 g/kg, i.p.; Sigma) at the acute injury stage (1.3 ± 0.5 days post-injury). After confirming the absence of the toe-pinch withdrawal reflex, the animal was placed in a supine position, and its body temperature was monitored and maintained at 37±1 °C using a temperature controller (model TC-1000; CWE Inc.). A tracheostomy was performed, and a PE-240 tube (Clay Adam) was inserted into the trachea below the larynx. The femoral artery was cannulated with a PE-50 tube for blood pressure measurement (DTX-1 transducer coupled with an amplifier; BPM-832; CWE Inc.) and blood gas analysis (iSTAT1 blood analyzer; Abbott). Then, the animal was placed in a prone position, followed by a dorsal incision and laminectomy at the third-fifth cervical vertebrae. Three oxygen sensor foils (3 × 3 mm^2^; SF-RPSu4; Presens) were placed on the dorsal cervical spinal cord with a 0.5 mm interval after durotomy (Fig. 1). One oxygen sensor foil was placed on the lesion epicenter (i.e., middle). The other two foils were placed rostral and caudal to the lesion epicenter. An oxygen detector unit (DU01; Presens) was placed above the cervical region with a focus on the oxygen sensor foils (Fig. 1). The peripheral oxygen saturation (SpO_2_) was monitored by a pulse oximeter clamped on the foot (PhysioSuite; Kent Scientific). The animal breathed a hyperoxic gas (50% O_2_ + 50% N_2_, 2 L/min) through a t-tube connected to the tracheal tube for 15 minutes, followed by 15 minutes of a normoxic gas (21% O_2_ + 79% N_2_, 2 L/min). The arterial blood gas was analyzed in the final minute of hyperoxic and normoxic breathing.

**Fig. 1.**
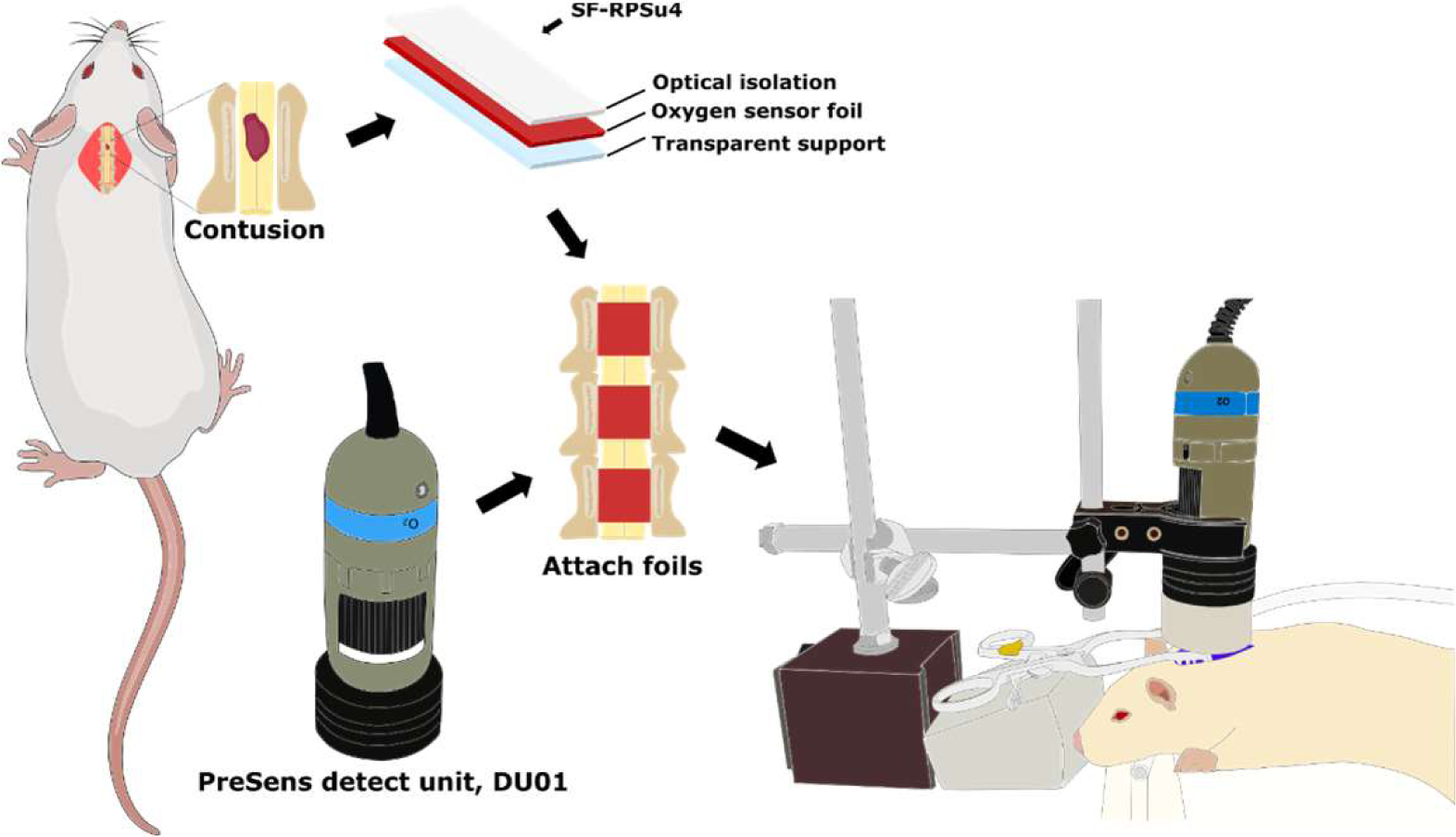
Schematic diagrams showing the experimental setup. The oxygen sensor foil comprises three layers: the optical isolation layer, the oxygen sensor layer, and the transparent support layer. Three oxygen sensor foils (3 × 3 mm^2^) were placed on the spinal cord after durotomy in the acute injury stage. One oxygen sensor foil was placed on the lesion epicenter, and the other two were placed on spinal segments rostral and caudal to the lesion site, respectively. The position of the detection unit was adjusted above the cervical spinal cord to image oxygen sensor foils.

### Data and statistical analyses

The blood pressure and SpO_2_ signals were digitized with CED Micro 1401-1 (Cambridge Electronic Design Ltd.) at a sampling rate of 1000 Hz using Spike 2 software (Cambridge Electronic Design Ltd.). The oxygen sensor foils were calibrated in a plastic chamber flushed with N_2_ (2 L/min) or normoxic gas (21% O_2_ + 79% N_2_, 2 L/min). The raw oxygen sensor foil image was converted into a calibrated image (Fig. 2A-B), and the oxygen sensor foil’s fluorescence was converted into a percentage air saturation value (% air sat.) using the VisiSens AnalytiCal 1 software (Presens). Figures 2C-D show that the oxygen level detected by the oxygen sensor foil correlated significantly with actual oxygen levels from 0% to 21%. During the experimental protocol, the oxygen sensor foils were imaged every minute after placement on the dorsal spinal cord.

**Fig. 2.**
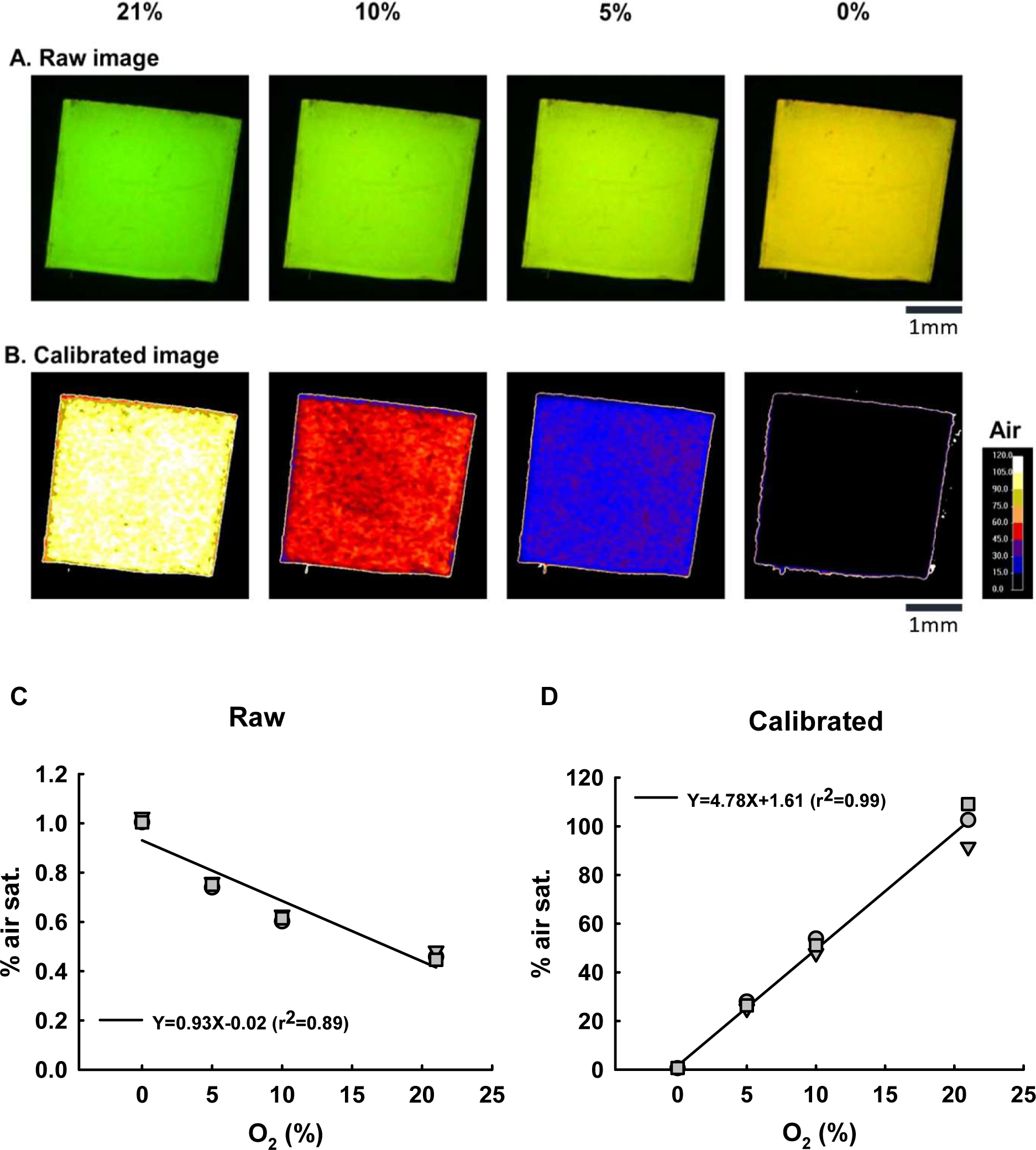
Calibration of the oxygen sensor foil under different oxygen concentrations. Panels A and B show the raw fluorescence and calibrated images under 21%, 10%, 5%, and 0% oxygen. Panels C and D show the raw and calibrated data (% air saturation [% air sat.]) under different oxygen levels. The results show that changes in the oxygen sensor foil’s fluorescence are highly correlated with the oxygen level, suggesting the feasibility of the oxygen sensor foil for oxygen detection. This correlation analysis was repeated three times using three oxygen sensor foils (circle, triangle, and square).

The systolic and diastolic blood pressure, heart rate, SpO_2_, and spinal cord oxygenation values were averaged over the final 3 minutes of hyperoxic and normoxic breathing. All parameters were analyzed using a two-way mixed-design analysis of variance (factor one: uninjured and contused; factor two: normoxia and hyperoxia) followed by the Student–Newman–Keuls post hoc test. All data are expressed as means ± standard deviations. A *P*-value <0.05 was considered statistically significant.

### Spinal cord histology

The animal was perfused with heparinized saline (250 mL), 4% paraformaldehyde (250 mL, Alfa Aesar), and 10% sucrose (J.T. Baker) in 4% paraformaldehyde (250 mL) after the completion of physiological recordings. The cervical spinal cord was dissected, placed in a phosphatase buffer solution containing 30% sucrose for cryoprotection, and cryosectioned into 40 µm slices using a cryostat (CM 1850; Leica). The spinal cord slices were stained with cresyl violet (Acros Organics) and imaged using a microscope (DM750; Leica) connected to a digital camera (EOS 600D; Canon). The spinal cord area was analyzed using ImageJ software (National Institutes of Health).

## Results

### Effect of hyperoxia on the spinal cord oxygenation

Cervical spinal cord contusion significantly impacted the spinal cord vascular structure, as shown by the shrinking of the dorsal spinal vessel at the lesion epicenter (Fig. 3). Spinal cord oxygenation could be detected using oxygen sensor foils attached to the dorsal spinal cord. The oxygen sensor foils fluoresced differently under normoxia and hyperoxia in both uninjured and contused rats, suggesting they could detect changes in spinal cord oxygenation (Fig. 3). The spinal cord oxygenation level was similar across the three spinal cord positions and could be maintained at ∼20%–23% (rostral: 22.7% ± 5.1%; middle: 20.3% ± 3.3%; caudal: 21.9% ± 4.2%) during normoxic breathing in uninjured animals. The spinal cord oxygenation level was significantly lower at all positions in contused animals (rostral: 15.5% ± 6.1%; middle: 14.8% ± 6.0%; caudal: 15.6% ± 5.0%) compared to uninjured animals (*P* < 0.05; Fig. 4A).

**Fig. 3.**
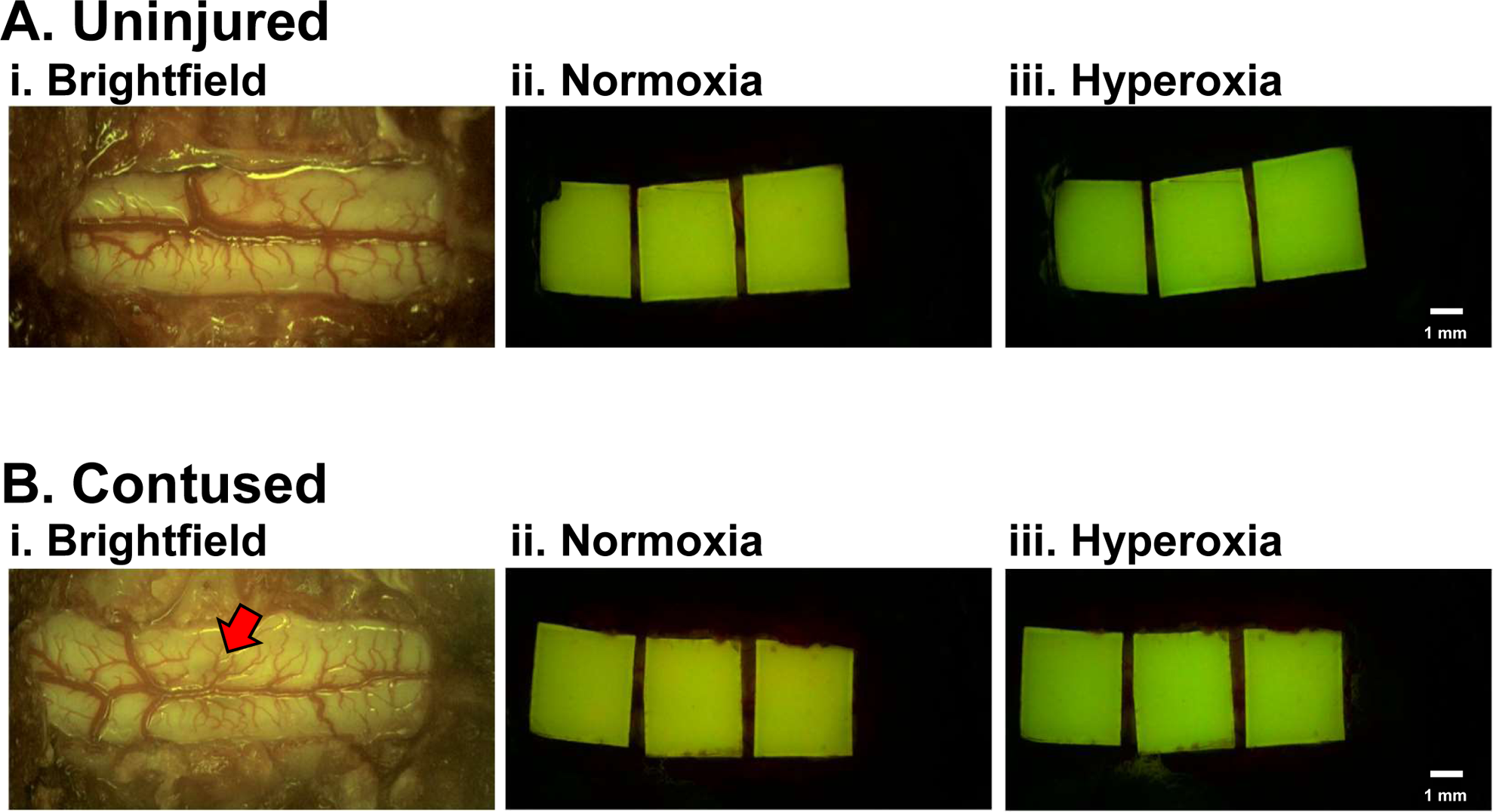
Representative images of oxygen sensor foils on the spinal cord of uninjured (A) and contused (B) animals. Panel i shows a brightfield image of the dorsal spinal cord. Panels ii and iii show fluorescent images of oxygen sensor foils placed on the spinal cord of an animal during normoxia (ii) or hyperoxia (iii) breathing. The color of oxygen sensor foils was visually different during normoxia (yellow) and hyperoxia (green). The red arrow in panel B (i) indicates the contused region.

**Fig. 4.**
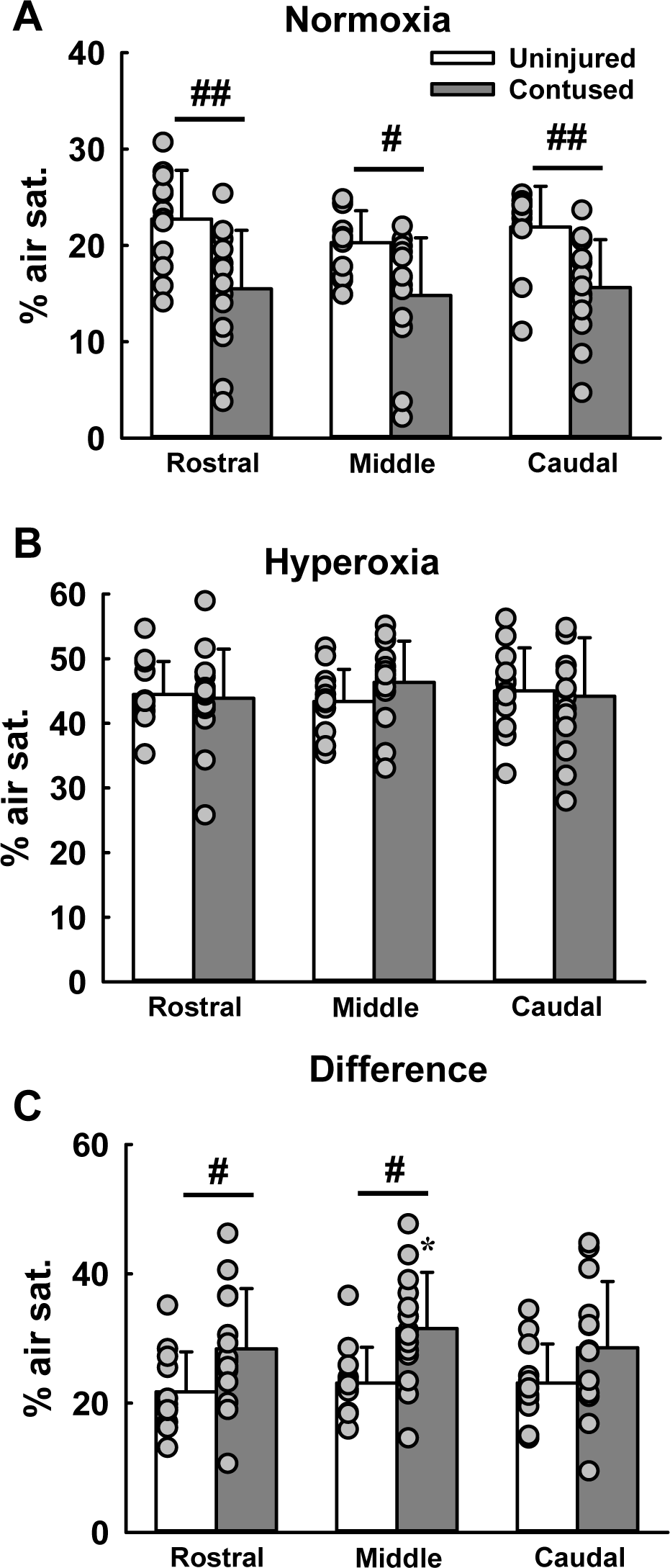
Spinal cord oxygenation during hyperoxia and normoxia. Panels A and B show the average spinal cord oxygenation levels in the rostral, middle, and caudal spinal cord segments compared to the lesion site under normoxia and hyperoxia. Panel C shows the change in spinal cord oxygen level from normoxia to hyperoxia. Data are presented as mean ± standard deviation (bar chart) and individual data points (grey circle). Sample size: uninjured (*n* = 12) and contused (*n* = 14).#, *P* < 0.05; ##, *P* < 0.01 between uninjured and contused animals; *, *P* < 0.05 vs. rostral and caudal spinal cord segments.

During hyperoxic breathing, the spinal cord oxygenation level increased significantly in both uninjured and contused animals (Fig. 4B). Specifically, the spinal cord oxygenation level increased to 44.5% ± 5.1%, 43.3% ± 5.0%, and 45.0% ± 6.7% at the rostral, middle, and caudal positions in uninjured rats, respectively (Fig. 4B). Similarly, hyperoxia could significantly increase the spinal cord oxygenation level of contused animals to ∼43%–47% (rostral: 43.9% ± 7.6%; middle: 46.3% ± 6.4%; caudal: 44.2% ± 9.1%), similar to the levels of uninjured animals (Fig. 4B). The capability of increasing spinal cord oxygenation between uninjured and contused animals was also calculated and shown in Figure 4C. The result showed that hyperoxia could increase spinal cord oxygenation by ∼21%–23% in uninjured rats (rostral: 21.7% ± 6.2%; middle: 23.1% ± 5.6%; caudal: 23.1% ± 6.1%). Notably, the increase in spinal cord oxygenation level at the rostral and middle positions was significantly greater in contused animals (28.4% ± 9.3% and 31.5% ± 8.7%, respectively) than in uninjured animals (*P* < 0.05; Fig. 4C). Moreover, the increase in spinal cord oxygenation level was significantly greater at the middle position than at the rostral and caudal positions in contused animals (*P* < 0.05; Fig. 4C).

### Effect of hyperoxia on the circulatory function

Representative examples of the blood pressure in uninjured and contused animals are shown in Figure 5. The systolic and mean arterial blood pressure were significantly lower in contused animals than in uninjured animals during normoxia (*P* < 0.05; Figs. 6A and 6C). Hyperoxia significantly elevated the blood pressure in both groups (Fig. 5). Specifically, mean arterial blood pressure increased from 57.0 ± 12.9 mmHg during normoxia to 81.7 ± 12.5 mmHg during hyperoxia in uninjured animals due to increases in both systolic and diastolic blood pressure (*P* < 0.01; Fig. 6A-6C). Similarly, mean arterial blood pressure significantly increased in contused animals during hyperoxia (normoxia: 40.5 ± 10.5 mmHg; hyperoxia: 62.9 ± 16.0 mmHg; Fig. 6C). However, the blood pressure parameters remained significantly lower in contused animals than in uninjured animals during hyperoxia (*P* < 0.05; Fig. 6A-6C). While cervical spinal cord injury significantly reduced blood pressure, heart rate was comparable between uninjured and contused groups, regardless of normoxia or hyperoxia (Fig. 6).

**Fig. 5.**
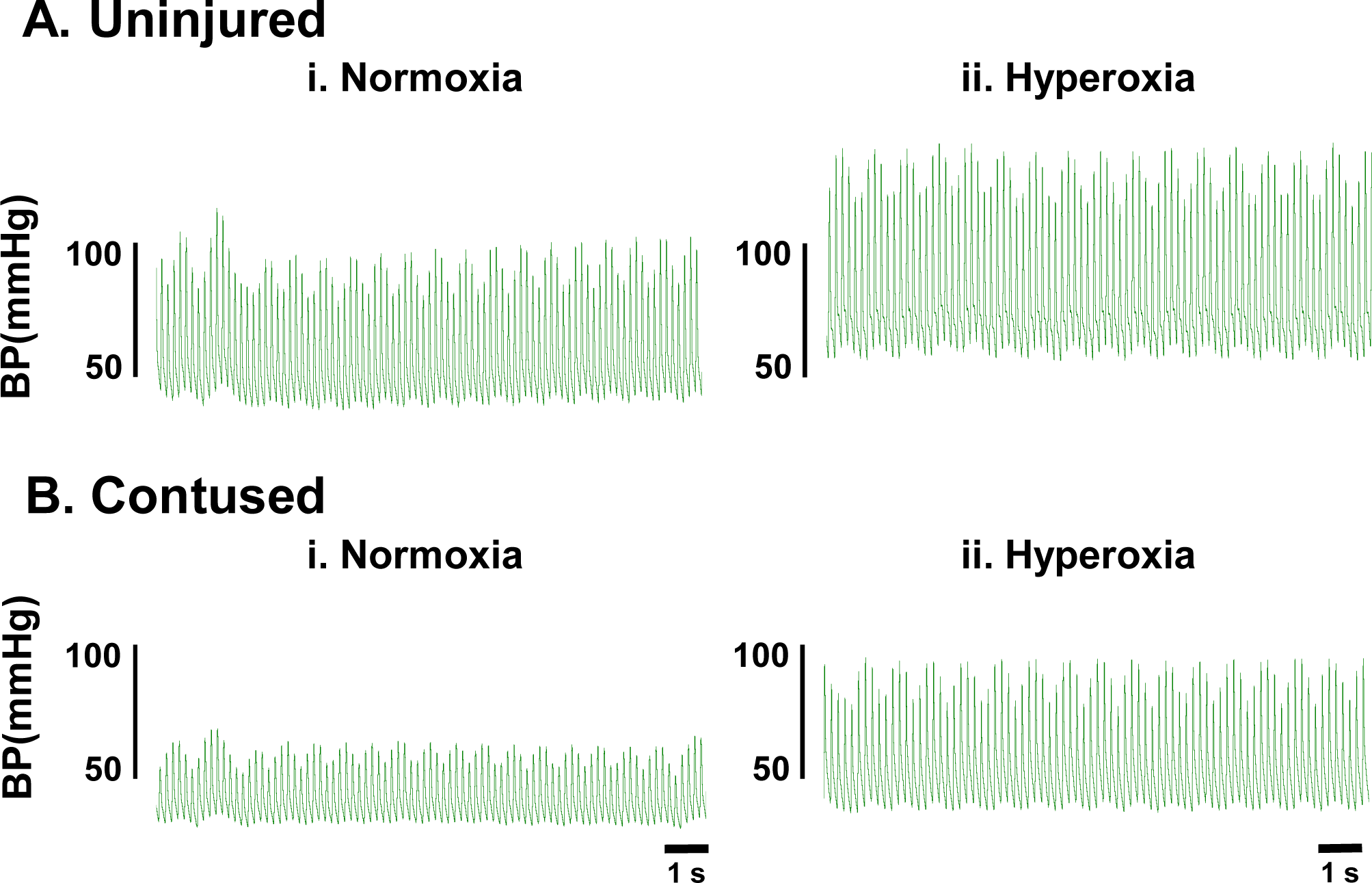
Representative examples of arterial blood pressure in an uninjured (A) and contused (B) animal during normoxia (i) and hyperoxia (ii) breathing. The blood pressure was lower in the contused than in the uninjured animal under both hyperoxic and normoxic conditions. Acute hyperoxia increased blood pressure in both uninjured and contused animals.

**Fig. 6.**
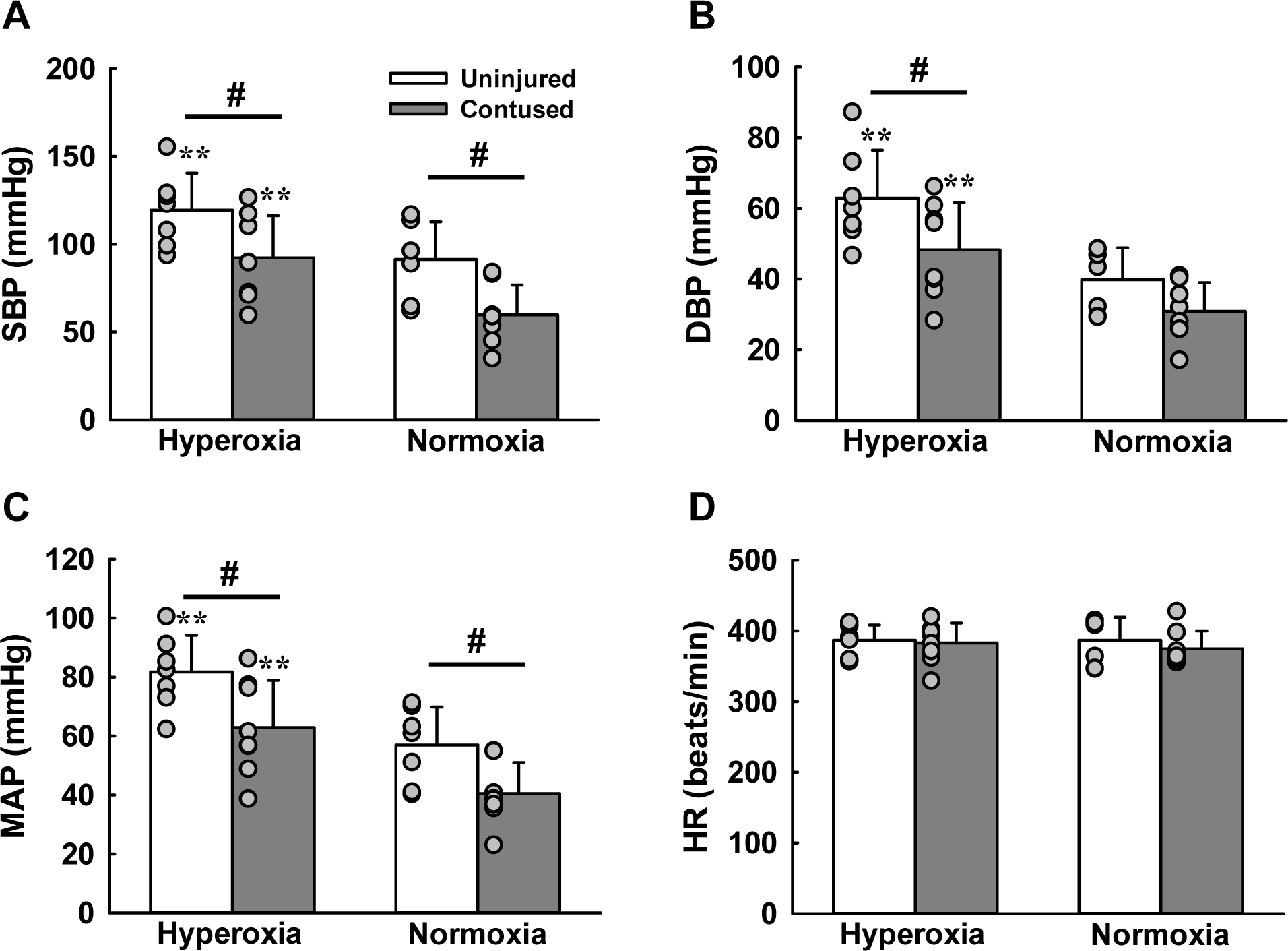
The averaged arterial blood pressure and heart rate of uninjured and contused animals during normoxic and hyperoxic breathing. The systolic blood pressure (SBP; A), diastolic blood pressure (DBP; B), and mean arterial blood pressure (MAP; C) were lower in contused animals than in uninjured animals. These blood pressure parameters increased during hyperoxic breathing. Heart rate (HR; D) was not significantly affected by cervical contusion or inhaled oxygen level. Data are presented as mean ± standard deviation (bar chart) and individual data points (grey circle). Sample size: uninjured (*n* = 7) and contused (*n* = 8). **, *P* < 0.01 between hyperoxia and normoxia; #, *P* < 0.05 between uninjured and contused animals.

### Effect of hyperoxia on the blood gas parameters and peripheral oxygen saturation

The blood gases were analyzed during normoxic and hyperoxic breathing in both groups. The arterial partial pressure of oxygen (PaO_2_) was significantly lower in contused (47.8 ± 6.2 mmHg) than in uninjured (63.0 ± 2.6 mmHg) animals (*P* < 0.01; Fig. 7A), resulting in significantly lower SpO_2_ in contused (71.3% ± 2.2%) than in uninjured (79.8% ± 4.5%) animals (*P* < 0.01; Fig. 7D). In addition, the arterial partial pressure of carbon dioxide (PaCO_2_) was higher in contused than in uninjured animals (*P* < 0.05, Fig. 7B).

**Fig. 7.**
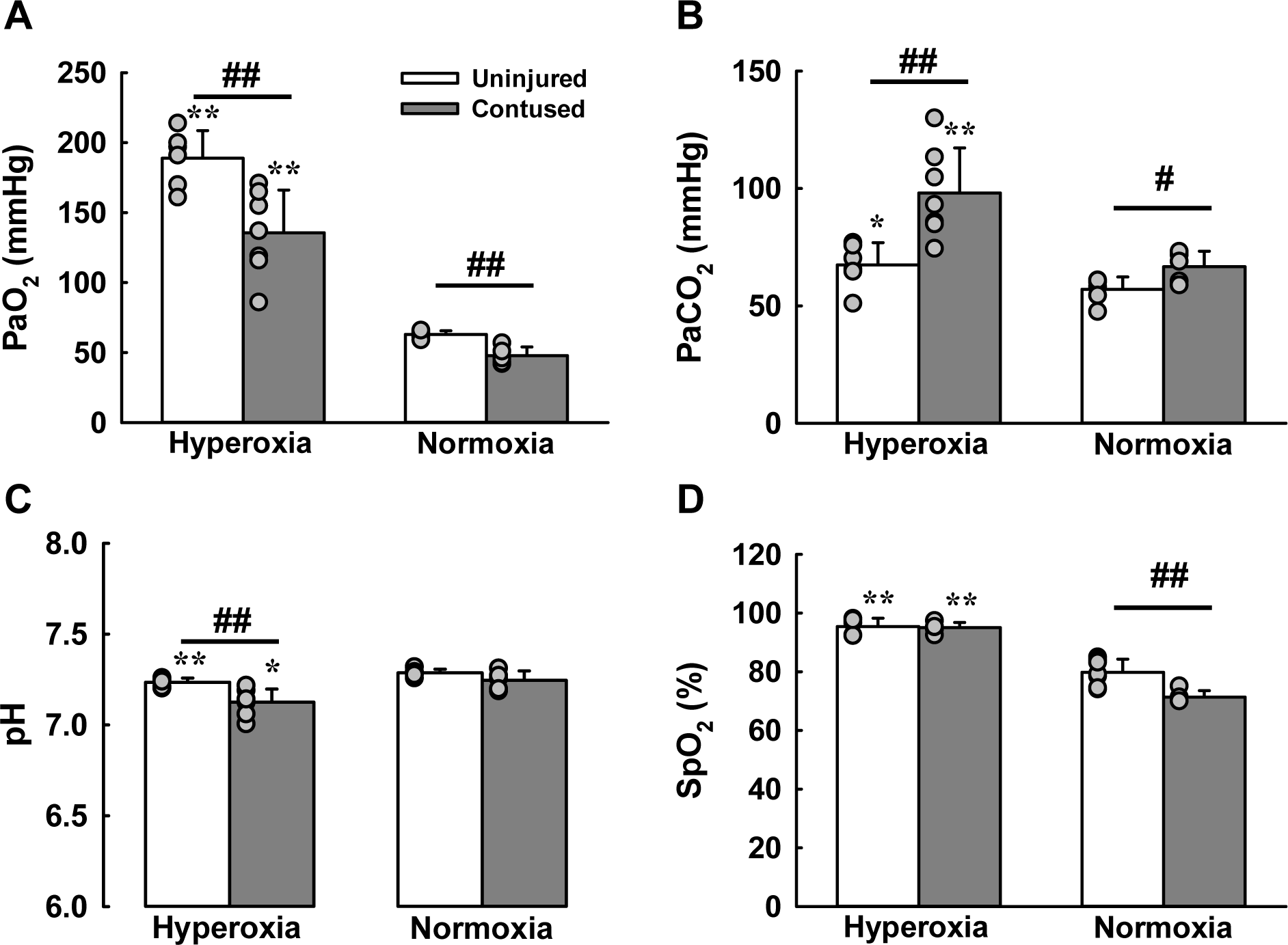
The average blood gas parameters and peripheral oxyhemoglobin saturation of uninjured and contused animals during normoxic and hyperoxic breathing. The arterial oxygen partial pressure (PaO_2_; A), arterial carbon dioxide partial pressure (PaCO_2_; B), and pH (C) were measured in arterial blood samples, and peripheral oxyhemoglobin saturation (SpO_2_, D) was monitored by pulse oximetry. During normoxic breathing, PaO_2_ was significantly lower and PaCO_2_ was significantly higher in contused than in uninjured animals. Hyperoxia significantly elevated PaO_2_ and PaCO_2_ in both groups, resulting in a significantly lower pH in contused animals. Data are presented as mean ± standard deviation (bar chart) and individual data points (grey circle). Sample size: uninjured (*n* = 6) and contused (*n* = 7). *, *P* < 0.05; **, *P* < 0.01 between hyperoxia and normoxia; #, *P* < 0.05; ##, *P* < 0.01 between uninjured and contused animals.

During hyperoxia, both PaO_2_ and PaCO_2_ significantly increased in both groups. Specifically, PaO_2_ increased to 188.8 ± 19.8 mmHg in uninjured animals and 135.6 ± 30.1 mmHg in contused animals (*P* < 0.01; Fig. 7A). Because the PaO_2_ of most animals was >100 mmHg during hyperoxic breathing, SpO_2_ was also significantly elevated and did not differ significantly between uninjured (95.4% ± 2.8%) and contused (95.0% ± 1.8%) animals (Fig. 7D). PaCO_2_ also significantly increased in uninjured animals from 57.1 ± 5.2 mmHg during normoxia to 67.4 ± 9.5 mmHg during hyperoxia (*P* < 0.05; Fig. 7B). Notably, the increase in PaCO_2_ was greater in contused animals, significantly increasing from 66.7 ± 6.6 mmHg during normoxia to 98.1 ± 19.2 mmHg during hyperoxia (*P* < 0.01; Fig. 7B). These increases in PaCO_2_ during hyperoxia reduced the pH in both groups (*P* < 0.05; Fig. 7C).

### Spinal cord section morphology

Representative examples of cervical spinal cord sections stained with cresyl violet are shown in Figure 8. Cervical spinal cord injury disrupted both grey and white matter tissue. Moreover, hemorrhage was observed in the grey matter, with significantly increased spinal cord areas in contused animals than in uninjured animals, suggesting contusion injury causes spinal edema.

**Fig. 8.**
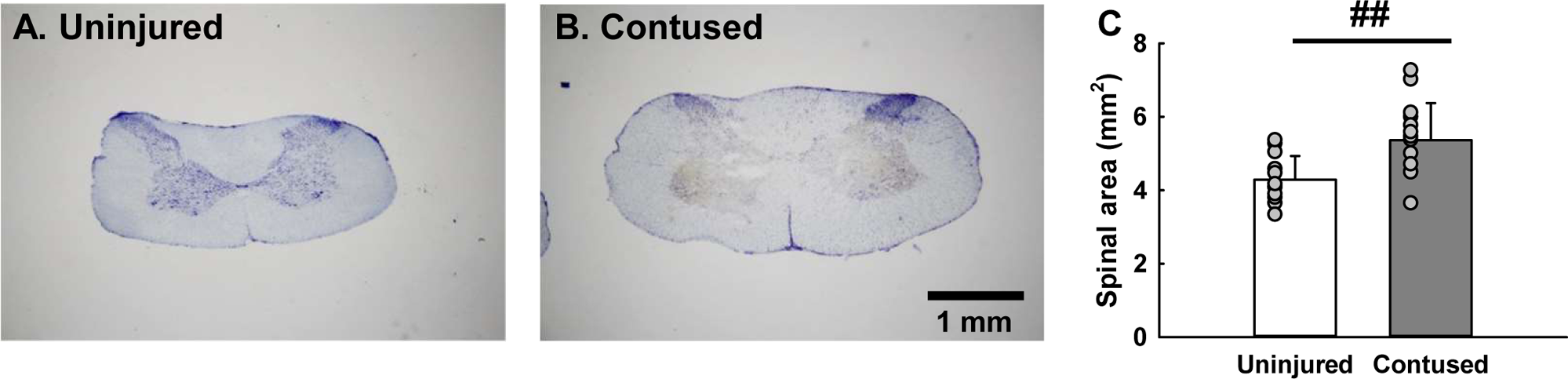
Representative examples of cervical spinal cord sections in an uninjured (A) and contused (B) animal. Panels A and B show a cervical spinal cord section stained with cresyl violet. Cervical spinal cord injury disrupts spinal cord grey and white matter. The contused animal also had a larger spinal cord (C), suggesting spinal cord edema was induced in the acute injury stage. Data are presented as mean ± standard deviation (bar chart) and individual data points (grey circle). Sample size: uninjured (*n* = 12) and contused (*n* = 14). ##, *P* < 0.01 between uninjured and contused animals.

## Discussion

### Methodological consideration

Tissue oxygenation can be measured using several techniques, including near-infrared spectroscopy, white light spectroscopy, optical/fluorescence-based oxygen monitoring, and reflectance spectroscopy imaging. These techniques have been used to monitor the oxygenation of the central nervous system in animal models (11, 16, 30, 37). However, these devices are expensive and may not be affordable in general labs. This study’s oxygen sensor foil and detector setup cost <3,000 USD. Its results show that the oxygen sensor foil can detect tissue oxygenation under *in vivo* conditions.

Two limitations of this method should be noted when interpreting our results. First, the oxygen sensor foil only detects oxygen near the dorsal spinal cord. Therefore, the spinal cord oxygenation level reflected the tissue oxygenation level in the dorsal portion of the spinal cord. Second, oxygen sensor foil fluorescence changes depend on interactions between oxygen and the sensitive layer, which needs several minutes to equilibrate. Therefore, the reaction time of the oxygen sensor foil is relatively longer than other techniques but remains feasible for monitoring tissue oxygenation under stable conditions.

### Impact of cervical spinal cord contusion on spinal cord oxygenation and circulatory function

This study showed that spinal cord oxygenation and circulatory function were significantly reduced after contusion injury. Several pathological conditions primarily caused the lower spinal cord oxygenation level. First, the cervical spinal cord injury interrupts the bulbospinal respiratory pathways and damages the phrenic nucleus within the cervical spinal cord, resulting in significant respiratory impairment and reduced oxygen supply to the central nervous system (17, 21, 25). Second, cervical spinal cord injury also damages the supraspinal vasomotor pathway innervating the sympathetic preganglionic neurons (1, 27). Therefore, sympathetic activity might be reduced, causing hypotension and hypoperfusion of the spinal cord (34, 35, 42). The impaired systemic and central hemodynamics attenuate the oxygen supply to the spinal cord. Third, the spinal cord vascular structure could be disrupted by contusion injury, reducing the blood flow supplying the spinal cord tissue around the injury epicenter (9, 22, 28).

### Therapeutic efficacy of hyperoxia on spinal cord oxygenation and circulatory function

Hyperoxia significantly elevated spinal cord oxygenation in both uninjured and contused groups. Notably, the increase in spinal cord oxygenation was greater in contused than in uninjured animals, leading to similar spinal cord oxygenation levels in contused and uninjured animals during hyperoxic breathing. In addition, the improvement in oxygenation was greater in the lesion’s epicenter than in the spinal regions rostral or caudal to it. These results suggest hyperoxia can effectively reverse the hypoxic spinal condition in contused rats.

Hyperoxia also significantly improved circulatory function, reflected in the increase in arterial blood pressure. The increases in systolic and diastolic blood pressure indicated that stroke volume and peripheral resistance increased during hyperoxic breathing. While some studies indicated that hyperoxia-induced vasoconstriction might reduce blood flow into the central nervous system and cause hypoxemia (2, 4), this was not observed in the current animal model of cervical spinal cord injury. Moreover, Dohi et al. reported that hypercapnia could induce vasodilation and increase spinal cord blood flow (6). Our current data showed that PaCO_2_ significantly increased during hyperoxic breathing. We speculate that hyperoxia-induced hypercapnia might induce vasodilation and counteract hyperoxia-induced vasoconstriction, maintaining or even increasing spinal cord blood flow and oxygen delivery.

### Clinical implications

Cervical spinal cord injury causes devastating pathological conditions in multiple physiological systems. Currently, there are no standard and effective treatments to restore physiological functions after cervical spinal cord injury. Hyperoxia is generally provided to patients under critical care and emergency conditions, and this approach is recommended for some disorders, including traumatic brain injury and chronic obstructive pulmonary disease (2). However, clinical and animal studies have shown that hyperoxia is associated with significant vasoconstriction and decreased cardiac output (31, 32), which might exacerbate existing pathological conditions. Our current results provide fundamental evidence showing that acute hyperoxia in the acute injury stage can improve spinal cord oxygenation and arterial blood pressure, suggesting this treatment could prevent spinal hypoxia and improve circulatory function. Future studies are needed to determine the optimal dosage and duration of normobaric hyperoxic treatment and evaluate whether it can improve functional recovery in the subchronic to chronic injury stages.

## Acknowledgements

The present study thanks Miss Rui-Yi Chen for preparing spinal cord histology.

